# Multitask group Lasso for Genome Wide association Studies in diverse populations

**DOI:** 10.1101/2021.08.02.454499

**Authors:** Asma Nouira, Chloé-Agathe Azencott

**Affiliations:** MINES ParisTech, PSL Research University, CBIO-Centre for Computational Biology, F-75006 Paris, France; Institut Curie, PSL Research University, F-75005 Paris, France INSERM, U900, F-75005 Paris, France

**Keywords:** Genome Wide Association Studies, Feature selection, Multitask group Lasso, Stability selection, Safe screening rules

## Abstract

Genome-Wide Association Studies, or GWAS, aim at finding Single Nucleotide Polymorphisms (SNPs) that are associated with a phenotype of interest. GWAS are known to suffer from the large dimensionality of the data with respect to the number of available samples. Other limiting factors include the dependency between SNPs, due to linkage disequilibrium (LD), and the need to account for population structure, that is to say, confounding due to genetic ancestry.

We propose an efficient approach for the multivariate analysis of multi-population GWAS data based on a multitask group Lasso formulation. Each task corresponds to a subpopulation of the data, and each group to an LD-block. This formulation alleviates the curse of dimensionality, and makes it possible to identify disease LD-blocks shared across populations/tasks, as well as some that are specific to one population/task. In addition, we use stability selection to increase the robustness of our approach. Finally, gap safe screening rules speed up computations enough that our method can run at a genome-wide scale.

To our knowledge, this is the first framework for GWAS on diverse populations combining feature selection at the LD-groups level, a multitask approach to address population structure, stability selection, and safe screening rules. We show that our approach outperforms state-of-the-art methods on both a simulated and a real-world cancer datasets.

## 1. Introduction

Over the last 15 years, Genome-Wide Association Studies (GWAS) have become one of the most prevalent methods to identify regions of the genome associated with complex phenotypic traits, and in particular complex diseases in humans.^1^ One of the major concerns in GWAS is population stratification, which arises when allele frequency differences between cases and controls are due to differences in genetic ancestry rather than to association with the phenotype. Many correction methods have been proposed to adjust the inflation of associations in diverse populations, including methods based on principal components analysis or on linear mixed models.^2^ However, it is possible that these techniques lead to overcorrection, in particular by masking population-specific disease loci.

An additional issue in GWAS is Linkage Disequilibrium (LD), which manifests as correlation between adjacent Single Nucleotide Polymorphisms (SNPs), creating statistical dependence between those markers and reducing statistical power.^3^ Combining strongly correlated SNPs into blocks, that is to say, groups of adjacent and correlated SNPs, and modeling the association signal over an entire region, is one way to address this limitation.

Classical approaches for GWAS are based on single-marker analyses, testing for association between each SNP and the phenotype independently. This may prevent the detection of effects that are due to SNPs acting additively, leading many authors to favor fitting a linear model to all SNPs jointly.^4^ Penalized regression approaches, such as the Lasso, which uses an ℓ_1_-norm regularization to shrink some coefficients of the model to zero, effectively removing them from the model, are particularly suited to this task.

Additional regularizers can be used to enforce additional prior hypotheses on the coefficients of such a linear model. Among them, the group Lasso^3,5^ ensures sparsity at the level of pre-defined groups of features, and the multitask Lasso^6,7^ fits models on related tasks jointly, encouraging similar sparsity patterns across all tasks.

In this work, we propose to combine both approaches into a multitask group Lasso framework, in which groups correspond to pre-defined LD patterns, and each task corresponds to a subpopulation, therefore simultaneously addressing the limitations of single-marker analyses and the issues of both LD and population structure.

In addition, we draw on the stability selection framework^8^ to improve the stability of the results, that is to say, their robustness to small perturbations in the input data, such as the removal of a few samples. Indeed, because the number of SNPs is typically much larger than that of samples, penalized regression approaches tend to select different sets of SNPs when presented with slightly different subsets of the same data, which severely limits their interpretability.

Finally, we use the recently proposed gap safe screening rules proposed by E. Ndiaye et al.^9^ to improve computational complexity, and scale our approach to about one million SNPs.

In what follows, we present our proposed approach in detail, place it in the context of existing work, and evaluate it on both a simulated data set and a real-world cancer GWAS data set.

## 2. Methods

Our proposed approach, MuGLasso, follows four steps, which we detail in this Section. First, we assign each sample to a genetic population, hence forming different but related tasks (Section 2.1). Second, we create LD-groups from correlations between SNPs, so as to perform feature selection at the level of groups rather than individual SNPs(Section 2.2). Third, we jointly fit one regularized model per task, using an *ℓ*_2,1_ penalty that enforces sparsity at the level of LD-groups (Section 2.3). Finally, we use stability selection to improve the robustness of the solution (Section 2.4).

### 2.1. Population stratification

Population structure, whereby the data is made of subsets of individuals that differ systematically both in genetic ancestry and in the phenotype under investigation, is a major confounding factor in GWAS. Indeed, it leads to detecting allele frequency differences in cases and controls that correspond to differences in ancestry, instead of a more direct association between genotype and phenotype. Several approaches have been developed to adjust for population structure.

Among them, a large number of methods rely on Principal Component Analysis (PCA),^10–12^ and consist of including top Principal Components (PCs) of the genotypes as covariates in regression models. In addition, linear mixed models^13^ can be used to model the phenotype as a combination of fixed and random effects, with the covariance of the latter being computed from a genetic similarity matrix. Although they often outperform PCA-based methods, the mixed model approaches tend to be more computationally demanding. Both approaches are similar in that regressing out principal components can be seen as approximation of a linear mixed model.^2^

However, these techniques may lead to ignoring population-specific SNPs, which is why we propose a multitask approach that can identify disease loci that are either population-specific or shared between populations. We therefore form tasks by separating the data into subpopulations, identified as clusters (using k-means clustering) on the projection of the genotypes on their top PCs.

### 2.2. Linkage disequilibrium groups

Linkage disequilibrium (LD) is the non-random association of alleles of at least two loci.^14^ LD can be leveraged to form groups of correlated SNPs. Grouping SNPs helps to alleviate the curse of dimensionality in GWAS by reducing the number of testing possibilities. This can be achieved by combining p-values within a group of correlated SNPs^15^, or through the use of penalized regression approaches that perform feature selection at the level of groups, rather than at the level of individual SNPs.^3^ The latter has the advantage over individual statistical testing of modeling the additive effects of multiple genetic markers simultaneously.

#### 2.2.1. Adjacency-constrained hierarchical clustering

In many species, including humans,^16^ LD is known to be correlated to the physical distance between SNPs. Hence, genomes can be clustered in LD blocks of strongly correlated adjacent SNPs, called in this paper LD-groups. Such LD-groups can be obtained using adjacency-constrained hierarchical agglomerative clustering,^3^ in which only physically adjacent clusters can be merged.

#### 2.2.2. LD-groups across populations

Because LD patterns may be influenced by genetic ancestry,^17^ we perform LD-groups partitioning for each population separately. We then combine those LD-groups into common shared LD-groups. More specifically, the set of coordinates of the boundaries of the shared LD-groups is obtained as the union of the sets of coordinates of the boundaries of the LD-groups for each population. This procedure is described in Supplementary Figure B1.

### 2.3. Multitask group Lasso

#### 2.3.1. General framework and problem formulation

We use a penalized regression approach to fit a multivariate linear model between the phenotype and the SNPs, with a regularization term that ensures that (1) the solution is sparse at the level of LD-groups and (2) the regression coefficients are smoothed within groups and across tasks. Such an approach provides shared LD-groups associated with the phenotype across all tasks, and allows for some LD-groups to be specific to each task.

##### Problem formulation

Given a set of *p* SNPs measured for *n* samples, we split the *n* samples in *T* subpopulations/tasks, each of size *n_t_* for *t* = 1,…, *T*, and the *p* SNPs in *G* LD-groups, each of size *p_g_* for *g* = 1,…,*G*. For each population *t*, we denote by 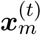 the *p*-dimensional vectors of SNPs of the *m*-th sample in the population (*m* = 1,…, *n_t_*), and by 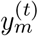 its phenotype. We then formulate the following optimization problem:

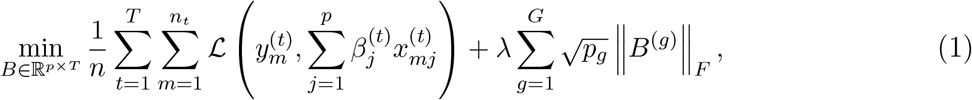

where *β*^(*t*)^ ∈ ℝ^*p*^ is the vector of regression coefficients specific to task *t* : *β*^(*t*)^ = (*B*_1*t*_,…, *B_pt_*), 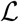 is the quadratic loss if the phenotype is quantitative (*y* ∈ ℝ) and the logistic loss if it is qualitative (*y* ∈ {0,1}), ||·||_*F*_ denotes the Frobenius norm, and *B*^(*g*)^ is a *p_g_* ×*T* matrix containing the regression coefficients, across all tasks, for the SNPs of group *g*.

#### 2.3.2. Related work

##### *ℓ*_2,1_-norm regularization

Our approach is closely related to the group Lasso^5^ and multitask Lasso,^6^ which both make use of an ℓ_2,1_-norm regularization. More precisely, the group Lasso corresponds to a special case of Equation (1), with a single task (*T* = 1), resulting in sparsity at the group levels. Using a group Lasso where the groups are defined based on LD blocks has been successfully applied to GWAS on up to 20 000 SNPs.^3^ The multitask Lasso corresponds to a special case of Equation (1), with each group containing exactly one SNP. This formulation ties sparsity patterns across tasks and has been applied before to multi-population GWAS, although only a few thousand SNPs.^7^

The multitask group Lasso we propose can also be reformulated as an ℓ_2,1_-norm regularization problem, through the creation of a new dataset 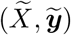 where 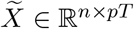 is a block-diagonal matrix such that each of the *T* diagonal blocks corresponds to the SNP matrix *X*^(*t*)^ ∈ ℝ^*n_t_*×*p*^ for task *t*, and 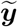 is a *n*-dimensional vector obtained by stacking the phenotype vectors for each task. Equation (1) can then be rewritten as:

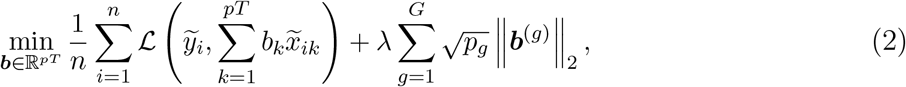

with ***b***^(*g*)^ ∈ ℝ^*p*_*g*_*T*^ the regression coefficients corresponding to all SNPs of group (*g*) for all tasks. In essence, this is a group Lasso with *G* groups each containing *T* copies (one per task) of the *p_g_* features of SNP group *g*. Thus *B_jt_* = ***b***_*p*_(*t*–1)_+*j*_.

##### Other multitask group Lassos

Other authors have proposed variations on the idea of a multitask group Lasso before. Several publications^18,19^ add a second regularization term to our formulation, increasing within-group or across-task sparsity. Unfortunately, this dramatically increases computational time, and indeed none of these publications analyze genome-wide data sets. In addition, because interpretation will be done at the group level rather than at the SNP level, within-group sparsity is not necessarily desirable in this context.

Several authors have built on these propositions and add a third regularization term, either enforcing group-independent task sparsity^20^ or overall sparsity (with an ℓ_1_-norm over all coefficients).^21^ Again, the addition of these regularizers severely hinders the applicability of these methods at a genome-wide scale due to computational limitations.

Hence none of these methods is readily applicable to our setting. In addition, their stability has never been evaluated, even though it is an important criterion for the reliability and interpretability of the results (see Section 2.4).

#### 2.3.3. Gap safe screening rules

To speed up the computation of the solution of Equation (2), we call upon gap safe screening rules,^9^ which are used to efficiently identify features for which the regression coefficients will be zero and hence ignore them when solving the problem. Such screening rules have been proposed for a large number of popular regularized regressions,^9^ including ℓ_2,1_-norm regularizations. In particular, Equation (2) can be solved using the `Gap_Safe_Rule` package^a^. We briefly summarize the idea underlying gap safe screening rules in Appendix B.3.

### 2.4. Stability selection

Unfortunately, in GWAS, penalized regression approaches often lack stability, that is to say, robustness to slight variations in the input dataset.^22^ However, stability increases both the reliability of the results and the interpretability. To address this limitation, *stability selection* ^8,22^ consists of performing feature selection repeatedly on subsamples of the data and only retains the features most often selected. More specifically, given a subsample *I* ⊆ {1,…, *n*} of size ⌊*n*/2⌋ of the data, we call 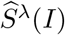 the set of features selected by the selection procedure of interest (for example, a Lasso), with hyperparameter λ, on this subsample of the data. For any feature *j* ∈ {1,…, *p*}, we call 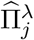 the probability that feature *j* is selected on a random subsample of size ⌊*n*/2⌋ of the data. This probability is determined, given *m* such random subsamples *I*_1_, *I*_2_,…, *I_m_*, as the proportion of those subsamples for which the feature selection procedure selects feature 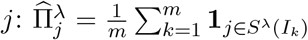. Finally, given a a threshold 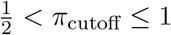 (in this work, we used π_cutoff_ = 0.75), the stable set of selected features is determined as 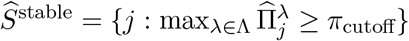.

## 3. Experiments

### 3.1. Data

#### Simulated data

Using GWAsimulator,^23^ we simulated GWAS data with realistic LD patterns from two populations (CEU : Utah residents with Northern and Western European ancestry and YRI: Yoruba in Ibadan, Nigeria) of the HapMap 3 data. We induced population structure by varying the case:control ratio within each subpopulation (CEU 1100:900 and YRI 900:1 100), as well as by simulating population-specific disease loci. We simulated a total of 149 970 disease SNPs, 2 999 (resp 4999) of which are specific to the CEU (resp. YRI) population (see Appendix A.1). The data contains 4000 samples and 1400 000 SNPs.

#### DRIVE Breast Cancer OncoArray

The DRIVE OncoArray dataset (dbGap study accession phs001265/GRU) contains 28281 individuals that were genotyped for 582 620 SNPs. 13 846 samples are cases and 14435 are controls. More details are available in Appendix A.2.

### 3.2. Preprocessing

#### Quality control and imputation

We removed SNPs with a minor allele frequency lower than 5%, a p-value for Hardy-Weinberg Equilibrium in controls lower than 10%, or a missing genotyping rate larger than 10%. We removed duplicate SNPs and excluded samples with more than 10% of SNPs missing. We imputed missing genotypes in DRIVE using IMPUTE2.^24^

#### LD pruning

We performed LD pruning using PLINK^26^ with a LD cutoff of *r*^2^ > 0.85 and a window size of 50Mb, both to reduce the number of SNPs and to better capture population structure using PCA.^25^ 1000000 SNPs remain in the simulated data and 313 237 in DRIVE.

#### PCA and population structure

We used PLINK^26^ to compute principal components of the genotypes. We thus identify two populations in the simulated data, matching the CEU and YRI populations (see Supplementary Figure C1a). In DRIVE, we identify two populations (see Supplementary Figure C1b), which we call POP1 (samples from the USA, Australia and Denmark) and POP2 (samples from the USA, Cameroon, Nigeria and Uganda).

#### LD-groups choice

We obtain LD-groups for each of the PCA-based populations using adjclust^27^ and obtain shared LD-groups as described in Section 2.2.2. Table 1 shows the number of LD-groups obtained for each subpopulation and the final number of shared groups.

**Table 1.** For each subpopulation of both datasets (simulated and real), LD-groups number is given and the shared LD-groups number after combination

| Data | Subpopulations | LD-groups number | Shared LD-groups number |
| --- | --- | --- | --- |
| Simulated data | CEU | 25 281 | 35 792 |
|  | YRI | 15 636 |  |
| DRIVE real data | POP1 | 8 152 | 17 782 |
|  | POP2 | 5 032 |  |

### 3.3. Comparison partners

As a baseline, we use PLINK^26^ to perform tests of association between each SNP and the phenotype, either using the top PCs as covariates (**Adjusted GWAS**), or treating each population separately (**Stratified GWAS**). We also compute a PCA-adjusted phenotype as the residuals of a regression between the top PCs and the phenotype. To evaluate the effects of grouping correlated SNPs and separating the populations in tasks, we compare MuGLasso to a Lasso (single task, no groups) on each population (**Stratified Lasso**) or on the adjusted phenotype (**Adjusted Lasso**), as well as a group Lasso (single task) on each population (**Stratified group Lasso**) or on the adjusted phenotype (**Adjusted group Lasso**).

For computational efficiency, we use bigLasso^28^ for the Lasso, and `Gap_Safe_Rule`^9^ for the group Lasso. For all methods, we set the regularization hyperparameter by cross-validation.

To compare these methods, we report runtime, ability to recover true causal SNPs (in the case of simulated data), and stability of the selection. To measure selection stability, we repeat the feature selection procedure on 10 subsamples of the data, and report the average Pearson’s correlation between all pairs of indicator vectors representing the selected features for each subsample (see Appendix B.4 for details).

## 4. Results

### 4.1. MuGLasso draws on both LD-groups and the multitask approach to recover disease SNPs

On the simulated data, we observe (Figure 1a) that MuGLasso is better than any other method at recovering the true disease SNPs. Performing feature selection at the level of LD-groups, rather than individual SNPs, improves performance. Indeed, the group Lassos and MuGLasso outperform the SNP-level Lassos. In addition, treating all samples simultaneously (as in MuGLasso or the adjusted approaches) also improves performance. This confirms our hypothesis that grouping features and using all samples simultaneously both alleviate the curse of dimensionality.

**Fig. 1.**
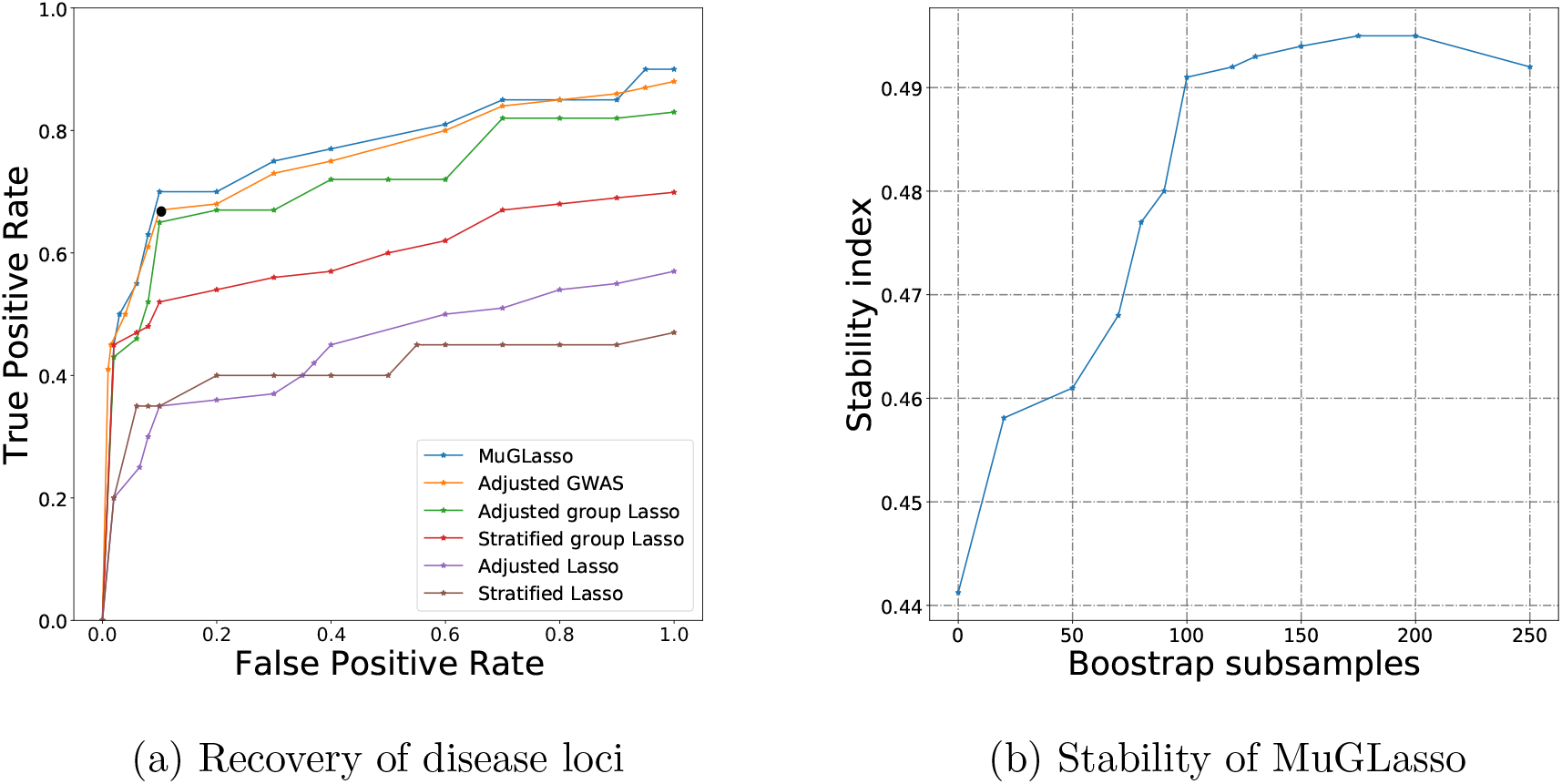
On simulated data, ability of different methods to retrieve causal disease SNPs as a ROC plot (1a), and stability index of MuGLasso as a function of the number of bootstrap samples (1b). On the ROC plot, the black dot indicates the performance of the stratified GWAS at the Bonferonni-corrected significance threshold.

On DRIVE, MuGLasso recovers 1051 SNPs in addition to all SNPs from the adjusted GWAS. They point to 32 risk genes that cannot be identified by the classical GWAS; half of those have been identified in meta-GWAS that included our samples, and another 7 have been associated with breast cancer risk or growth in other studies (see Supplementary Table C1).

However, this increased ability to recover relevant SNPs comes with an increase in computational time (see Supplementary Figure C2 on simulated data and Figure 2a on DRIVE). However, the implementation is efficient enough to allow computations on 10^6^ SNPs, even with the added cost of repeated subsampling to increase stability.

**Fig. 2.**
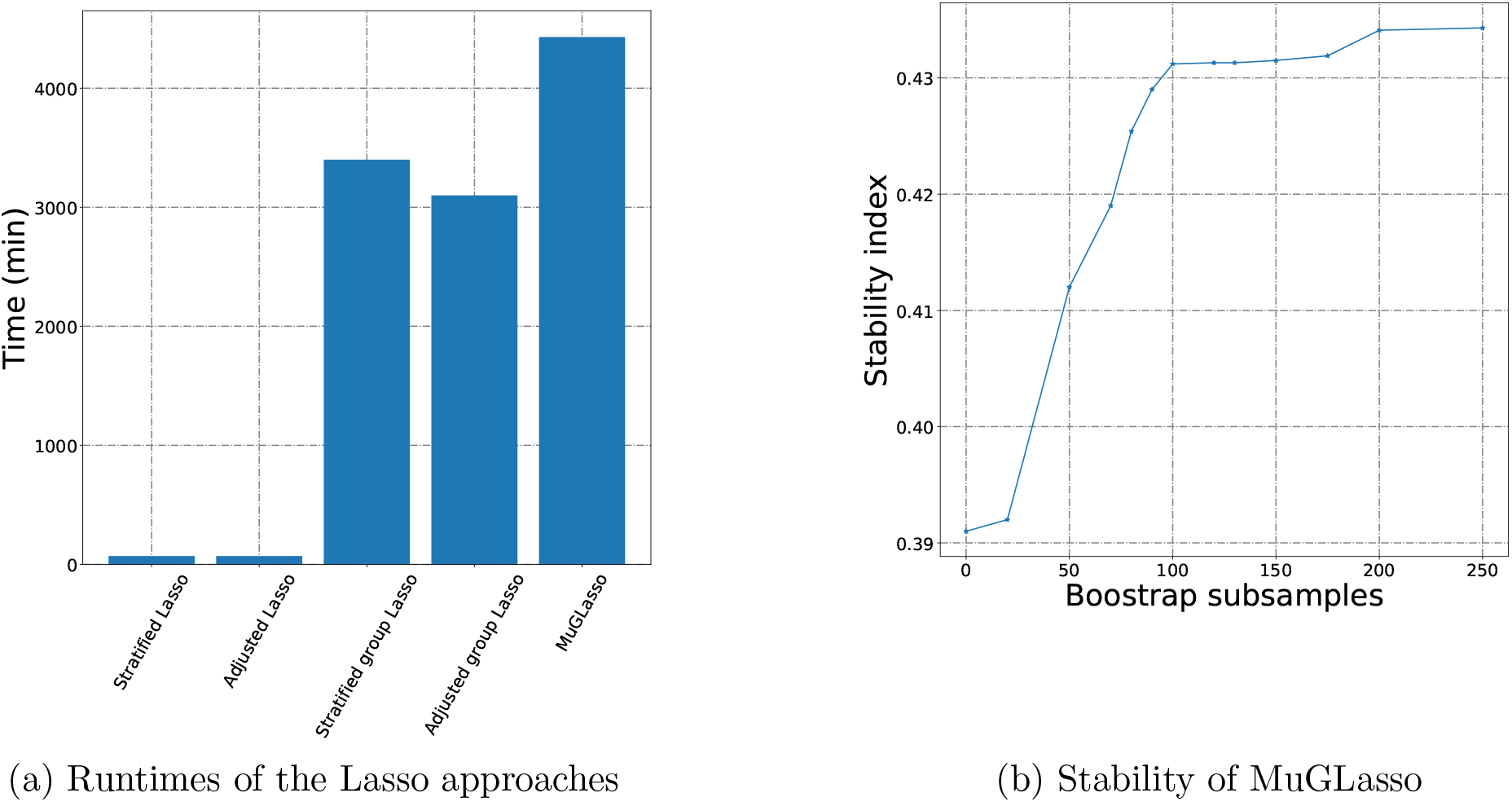
On DRIVE, runtimes of the different Lasso approaches (2a) and stability index of MuGLasso as a function of the number of bootstrap samples (2b).

### 4.2. MuGLasso provides the most stable selection

Figures 1b (simulated data) and 2b (DRIVE) show the stability index of MuGLasso as a function of the number of subsamples. Increasing the number of subsamples increases the stability of the selection. We use 100 bootstrap samples in all subsequent experiments as it appears to be an acceptable trade-off between runtime and stability.

Tables 2 and 3 show the stability index of the different methods, on simulated data and DRIVE, respectively. We ran the adjusted GWAS once on the entire data set, as would usually be done, and therefore cannot report its stability. Our results again illustrate that stability selection does increase the stability of Lasso methods. We confirm this by running MuGLasso without stability selection as well as Adjusted group Lasso with stability selection on top. In both cases, the stability index increases when stability selection is used. In addition, we report the total number of selected SNPs and LD-groups. For methods that select individual SNPs, we obtain the number of selected LD-groups by considering that each selected SNP selects its entire LD-group. Our results illustrate that the improved stability of MuGLasso does not come at the expense of selecting more features. On the contrary, stability selection provides fewer SNPs/LD-groups with better stability.

### 4.3. MuGLasso selects both task-specific and global LD-groups

For both datasets, the LD-groups selected by MuGLasso are a mixture between population-specific LD-groups (identified as those with near-zero regression coefficients for one task) and LD-groups that are shared between both populations. Table 4 shows the number of LD-groups/SNPs in each of these categories for MuGLasso. By contrast, the adjusted approaches do not provide population-specific LD-groups or SNPs.

Finally, we report on Figure 3 the precision and recall of MuGLasso and the stratified approaches on the population-specific SNPs. MuGLasso outperforms all other approaches in both precision and recall.

**Fig. 3.**
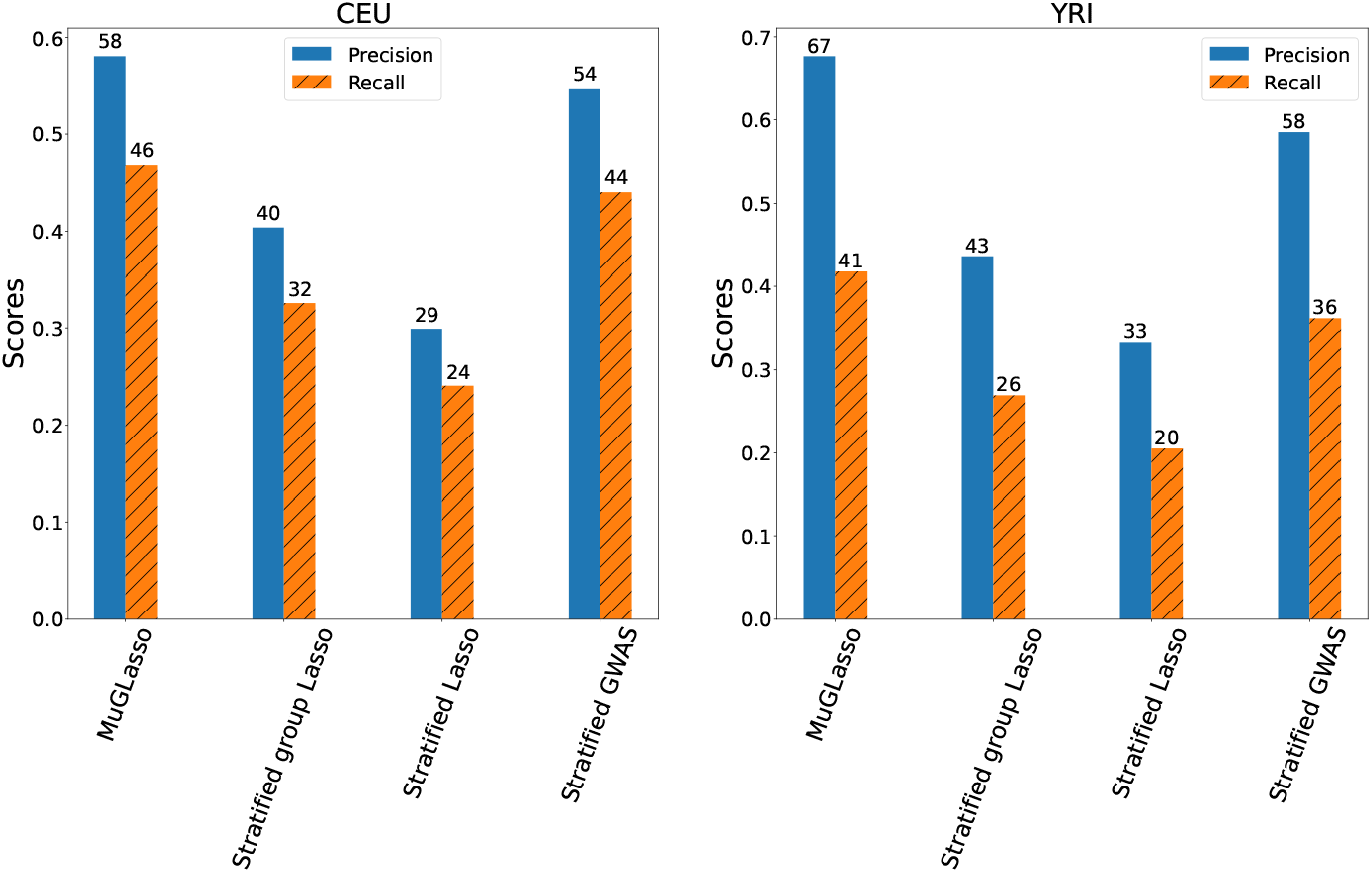
For simulated data, precision and recall of MuGLasso and the stratified approaches on the populations-specific SNPs

**Table 2.** Stability index and number of selected features for different methods, on simulated data

| Methods | Number of selected LD-groups | Number of selected SNPs | Stability index | Selection level |
| --- | --- | --- | --- | --- |
| MuGLasso | 5 623 | 155 312 | 0.4912 | LD-groups |
| MuGLasso without stab sel | 6 124 | 161 221 | 0.4412 | LD-groups |
| Adjusted group Lasso + stab sel | 6 054 | 162 104 | 0.4134 | LD-groups |
| Adjusted group Lasso | 6 347 | 167 204 | 0.3714 | LD-groups |
| Stratified group Lasso | 4 836 | 154 732 | 0.3398 | LD-groups |
| Adjusted Lasso | 5 379 | 158 856 | 0.2368 | Single-SNP |
| Stratified Lasso | 5 704 | 168 158 | 0.1742 | Single-SNP |
| Adjusted GWAS | 5 063 | 141 340 | - | Single-SNP |

## 5. Discussion and Conclusions

We presented MuGLasso, an efficient approach for detecting disease loci in GWAS data from diverse populations. Our approach is based on a multitask framework, where input tasks correspond to subpopulations, and feature selection is performed at the level of LD-groups. Assigning samples from PCA-identified populations to different tasks addresses the issue of population stratification, and retains the flexibility of identifying population-specific disease loci. Treating all samples together, by contrast with stratified approaches, alleviates the curse of dimensionality. Ensuring sparsity at the level of LD-groups addresses the high correlation between nearby SNPs and also alleviates the curse of dimensionality. Although more time-consuming than a classical GWAS, our implementation is computationally efficient enough to scale to the analysis of entire GWAS data sets of about one million SNPs.

**Table 3.** Stability index and number of selected features for different methods, on DRIVE

| Methods | Number of selected LD-groups | Number of selected SNPs | Stability index | Selection level |
| --- | --- | --- | --- | --- |
| MuGLasso | 62 | 1 357 | 0.4312 | LD-groups |
| MuGLasso without stab sel | 72 | 1 524 | 0.3911 | LD-groups |
| Adjusted group Lasso + stab sel | 59 | 1 293 | 0.3234 | LD-groups |
| Adjusted group Lasso | 68 | 1 466 | 0.2613 | LD-groups |
| Stratified group Lasso | 58 | 1 119 | 0.2498 | LD-groups |
| Adjusted Lasso | 41 | 874 | 0.2068 | Single-SNP |
| Stratified Lasso | 38 | 789 | 0.1581 | Single-SNP |
| Adjusted GWAS | 16 | 306 | - | Single-SNP |

**Table 4.**
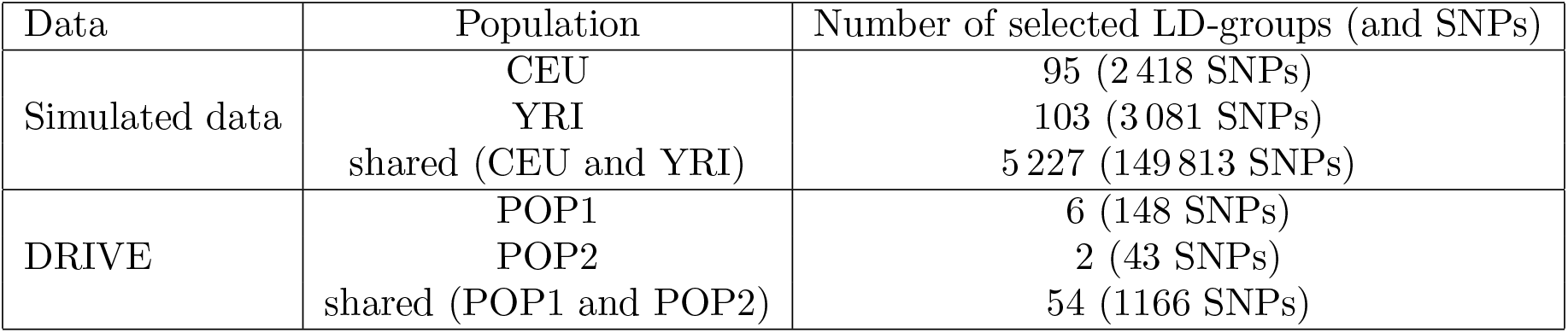
For MuGLasso, number of selected LD-groups/SNPs, across and per population

On simulated data, MuGLasso outperforms state-of-the-art approaches in its ability to recover disease loci. This also holds for population-specific SNPs; hence performance is not driven solely by the ability to recover disease loci that are common to all populations. In addition, MuGLasso is the most stable of all evaluated method, which increases interpretability.

Finally, although we presented MuGLasso in the context of admixed populations, our tool could be used in other multitask settings. In particular, tasks can stem from related phenotypes^18^ or from different studies pertaining to the same trait, in a meta-analysis approach.^19^ Groups could also be defined according to different prior biological knowledge, for example based on functional units such as genes, in the spirit of gene-set analyses of GWAS data. In addition, although we only presented results on case-control studies with two populations, the method directly applies to quantitative phenotypes and any number of tasks.

An important outcome of our study is that, although we have not included in MuGLasso a regularization term that would enforce sparsity at the level of tasks as in Li et al. (2020),^21^ we still obtain task-specific groups. Including such an additional term in Equation (1) would perhaps improve the already state-of-the-art task-specific precision and recall of MuGLasso, but this would unfortunately come at the expense of a notable increase in computational time, if only because of the cross-validation needed to set the value of a second hyperparameter.

An in-depth biological analysis of the loci identified by MuGLasso on DRIVE would illustrate the biological relevance of our method, but is out of the scope of this methodological paper.

In the future, we are looking forward to making use of the post-inference selection framework for group-sparse linear models^29^ to provide p-values for the selected loci. As of now, it is unclear how to apply these ideas to case-control studies in a computationally efficient manner.

## Acknowledgments

This work was supported by the French Agence Nationale de la Recherche (ANR-18-CE45-0021-01, ANR-19-CE14-0015 and ANR-19-P3IA-0001). OncoArray genotyping and phenotype data harmonization for the Discovery, Biology, and Risk of Inherited Variants in Breast Cancer (DRIVE) breast-cancer case control samples was supported by X01 HG007491 and U19 CA148065 and by Cancer Research UK (C1287/A16563). We thank Héctor Climente-González, Vivien Goepp and Lotfi Slim for discussions, and acknowledge the contributions of the following reviewers: Carly Bobak, Meghan Muse, Lauren McDonnell, Amy Farnham, Michael Mariani, Andrew Hederman and Ariana Haidari.

## Supplementary Materials and code

Supplementary materials are available online at https://www.biorxiv.org/content/10.1101/2021.08.02.454499. Code is available at https://github.com/asmanouira/MuGLasso_GWAS.

## Appendix A. Data availability

### Appendix A.1. Simulated data

Code to reproduce our simulations is available on https://github.com/asmanouira/MuGLasso_GWAS

Table A1 shows the location of the predefined disease loci, for each population. Table A2 shows the number of predefined disease loci, both common to both population and specific to each population.

**Table A1.** For simulated data, location of pre-defined disease loci represented by start/end positions information in each subpopulation through chromosomes: 2, 12, 19, 21 and 22.

| Chromosome | Subpopulations |  |
| --- | --- | --- |
|  | CEU | YRI |
| 2 | 1 000 - 50 000 | 1 000 - 50 000 |
| 12 | 10 - 37 000 | 10 - 40 000 |
| 19 | 1 000 - 50 000 | 1 000 - 50 000 |
| 21 | 10 - 10 000 | 10 - 7 000 |
| 22 | - | 10 - 2 000 |

**Table A2.**
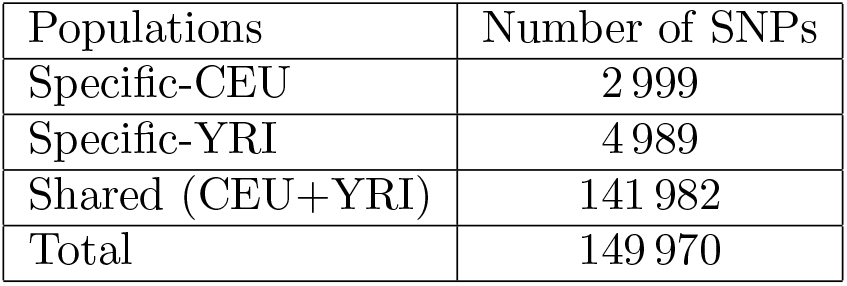
For simulated data, number of predefined causal SNPs

| Populations | Number of SNPs |
| --- | --- |
| Specific-CEU | 2 999 |
| Specific-YRI | 4 989 |
| Shared (CEU+YRI) | 141 982 |
| Total | 149 970 |

### Appendix A.2. DRIVE

#### Data access

The dataset “General Research Use” in DRIVE Breast Cancer OncoArray Genotypes is available from the dbGaP controlled-access portal, under Study Accession phs001265.v1.p1 (https://www.ncbi.nlm.nih.gov/projects/gap/cgi-bin/study.cgi?study\_id=phs001265.v1.p1). Researchers can gain access the data by applying to the data access committee, see https://dbgap.ncbi.nlm.nih.gov.

#### Ethics approval

The dataset was obtained from NIH after ethical review of project #17707, titled ”Network-guided multi-locus biomarker discovery”, and used under approval of this request (#67806-4).

## Acknowledgments

OncoArray genotyping and phenotype data harmonization for the Discovery, Biology, and Risk of Inherited Variants in Breast Cancer (DRIVE) breast-cancer case control samples was supported by X01 HG007491 and U19 CA148065 and by Cancer Research UK (C1287/A16563). Genotyping was conducted by the Center for Inherited Disease Research (CIDR), Centre for Cancer Genetic Epidemiology, University of Cambridge, and theNational Cancer Institute. The following studies contributed germline DNA from breast cancer cases and controls: the Two Sister Study (2SISTER), Breast Oncology Galicia Network (BREOGAN), Copenhagen GeneralPopulation Study (CGPS), Cancer Prevention Study 2 (CPSII), The European Prospective Investigation intoCancer and Nutrition (EPIC), Melbourne Collaborative Cohort Study (MCCS), Multiethnic Cohort (MEC), NashvilleBreast Health Study (NBHS), Nurses Health Study (NHS), Nurses Health Study 2 (NHS2), Polish Breast CancerStudy (PBCS), Prostate Lung Colorectal and Ovarian Cancer Screening Trial (PLCO), Studies of Epidemiologyand Risk Factors in Cancer Heredity (SEARCH), The Sister Study (SISTER), Swedish Mammographic Cohort (SMC), Women of African Ancestry Breast Cancer Study (WAABCS), Women’s Health Initiative (WHI).

## Appendix B. Supplementary Methods

### Appendix B.1. LD groups across populations

Figure B1 illustrates the process by which we obtain LD-groups across populations, from LD-groups obtained on each population separately using adjacency-constrained hierarchical clustering (see Section 2.2.1)

**Fig. B1.**
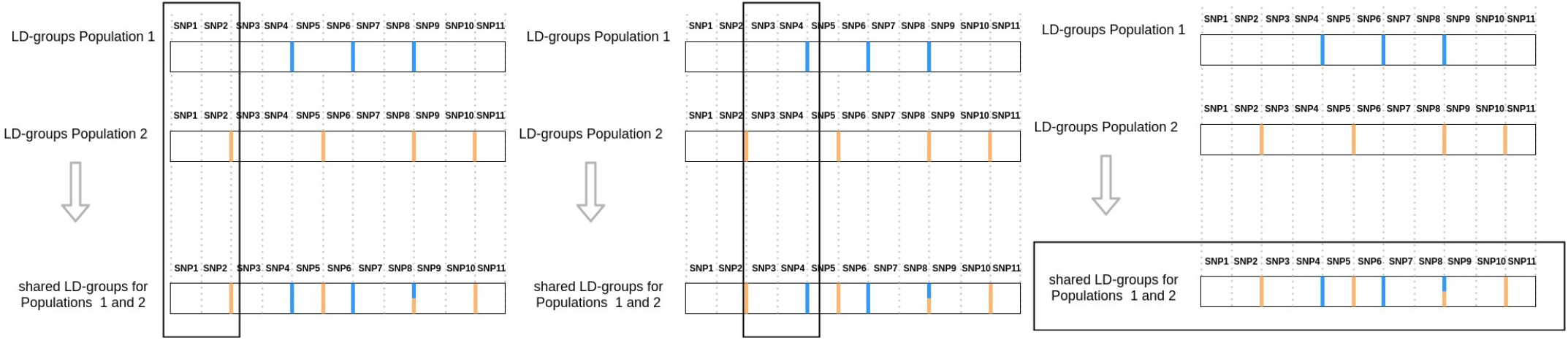
Choice of shared LD-groups choice after adjacency-constrained hierarchical clustering for each population

### Appendix B.2. Multitask group lasso

Figure B2 illustrates the architecture of the multitask group Lasso described in Section 2.3.

**Fig. B2.**
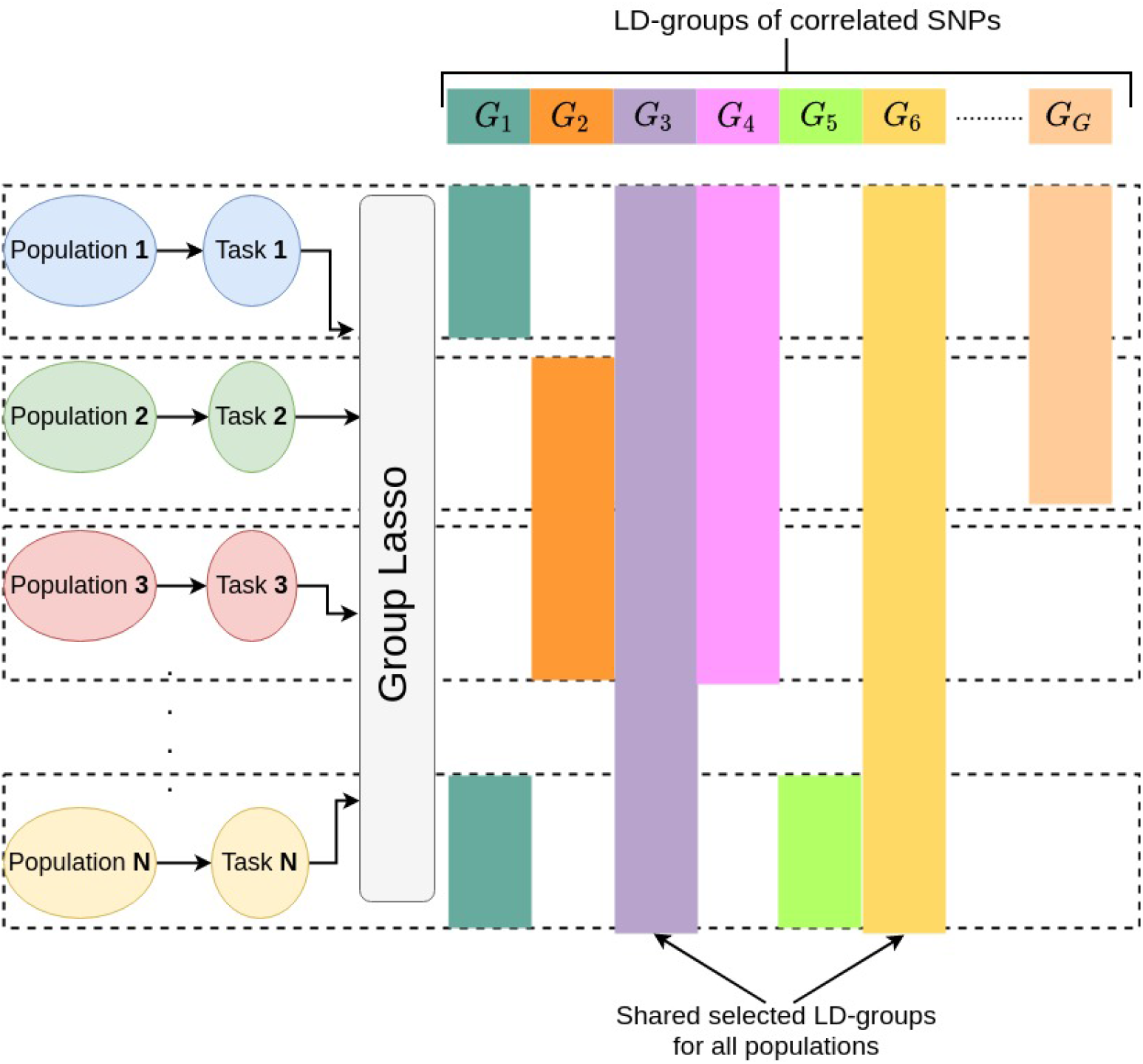
Multitask group Lasso architecture

### Appendix B.3. Gap safe screening rules

Let *X* ∈ ℝ^*n*×*d*^ be a design matrix and *y* ∈ ℓ^*n*^ the corresponding vector of outcomes, which can be binary or real-valued. We consider the following optimization problem:

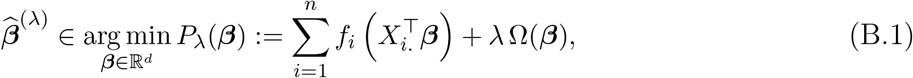

where all *f_i_* : ℝ → ℝ are convex and differentiable functions with 1/*γ*–Lipschitz gradient, and Ω : ℝ^*d*^ → ℝ_+_ is a norm that is group-decomposable, i.e., the set of *d* features is partitioned in *G* groups of sizes *d*_1_, *d*_2_,…, *d_G_*, and

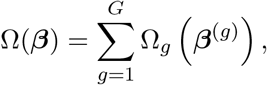

where each Ω_*g*_ is a norm on ℝ^*dg*^ and, as previously, ***β***^(*g*)^ corresponds to the coefficients of *β* restricted to the features in group *g*. As before, the λ parameter is a non-negative constant controlling the trade-off between the data fitting term and the regularization term.

Equation (2) is a special case of Equation (B.1) because the squared loss and the logistic loss are convex and differentiable.

Safe screening rules make it possible to solve such problems more efficiently by discarding features whose coefficients are guaranteed to be zero at the optimum, prior to using a solver. They usual rely on the dual formulation of Equation (B.1):

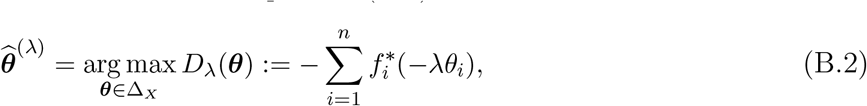

where 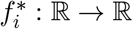 is the Fenchel-Legendre transform of *f_i_*, defined by 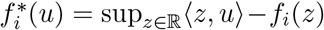 and Δ_*X*_ ⊂ ℝ^*n*^ is defined by 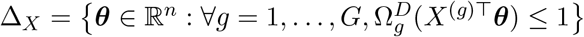, where 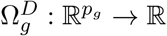 is the conjugate norm of Ω_*g*_, defined by 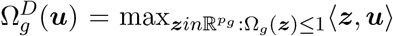, and *X*^*g*^ ∈ ℝ^*n*×*p*_*g*_^ is the design matrix *X* restricted to the features/columns in group *g*.

In our setting,

- 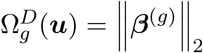 and 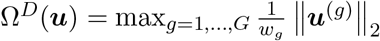.
- If one uses the squared loss, that is to say, 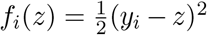, then 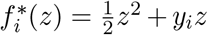 and the Lipschitz constant is *γ* = 1.
- If one uses the logistic loss, that is to say, ***y*** ∈ {0,1}^*n*^ and *f_i_*(*z*) = −*y_i_z* + log(1 + exp(*z*)), then

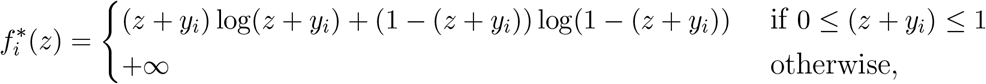

and the Lipschitz constant is *γ* = 4.

The general idea of safe screening rules, introduced by [EGVR10], is to find a region 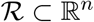 such that if 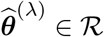, for any *g* ∈ {1,…, *G*},

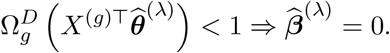

Gap safe screening rules [N^+^17] exploit the duality gap (*P*_**λ**_(***β***) – *D*_**λ**_(***θ***)) to obtain the radius of the safe region 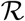. More specifically, Ndiaye et al. show that ∇***β*** ∈ ℝ^*p*^, ∇***θ*** ∈ Δ_*X*_,

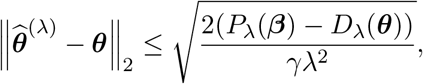

which leads them to define, for any ***β*** ∈ ℝ^*p*^ and ***θ*** ∈ Δ_*X*_, the ball centered in ***θ*** and of radius 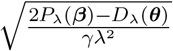 as a safe region, that is to say a region that is guaranteed to contain 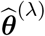.

### Appendix B.4. Measuring selection stability

To measure the stability of a feature selection property, we use the sample’s Pearson coefficient [NB16]. This stability index is closely related to that proposed by Kuncheva [Kun08] and is appropriate for the comparison of feature sets of different sizes. This index relies on repeating the feature selection procedure *M* time (in this work, *M* = 10) and evaluating the overlap if the *M* resulting feature sets.

Each of the M sets of selected features can be represented by an indicator vector *s* ∈ {0,1}^*p*^, where *s_j_* = 1 if feature j is selected and 0 otherwise. The stability index between two feature sets 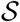 and 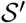, represented by their indicator vectors **s** and **s**’, is computed as the Pearsons’s correlation between these two vectors:

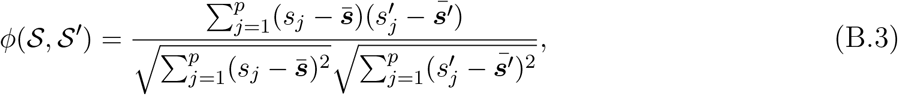

where 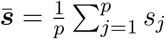 and 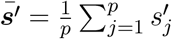.

Note that, because 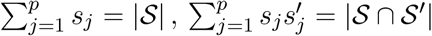, and 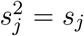, we can rewrite Equation (B.3) as

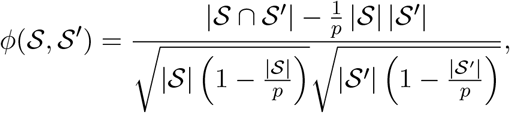

hence interpreting this index as the size of the intersection of the two sets, corrected by chance, that is to say, ensuring that the expected value of the index is 0 when the two selections are random.

The stability index between *M* sets of selected features is computed as the average pairwise stability index between all possible pairs of sets of selected features:

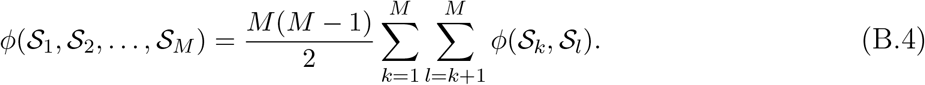

### Appendix C. Supplementary Results

#### Appendix C.1. PCA of the genotypes

Figure C1 shows the genotypes of the simulated data (Figure C1a) and the DRIVE data (Figure C1b) projected on the two first principal components of the data.

#### Appendix C.2. Runtimes

Figure C2 shows the runtimes of the different Lasso methods on simulated data.

**Fig. C1.**
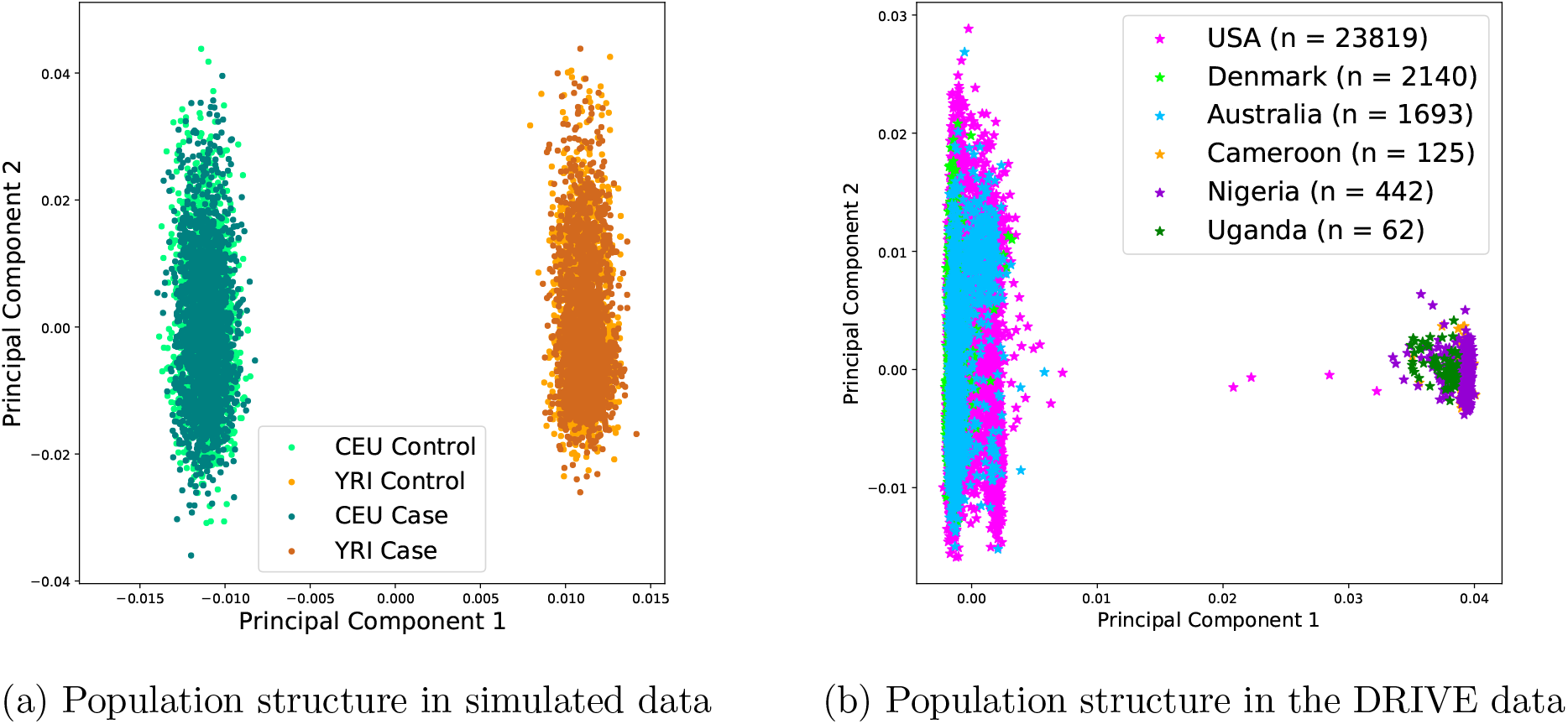
PCA for simulated and real datasets

**Fig. C2.**
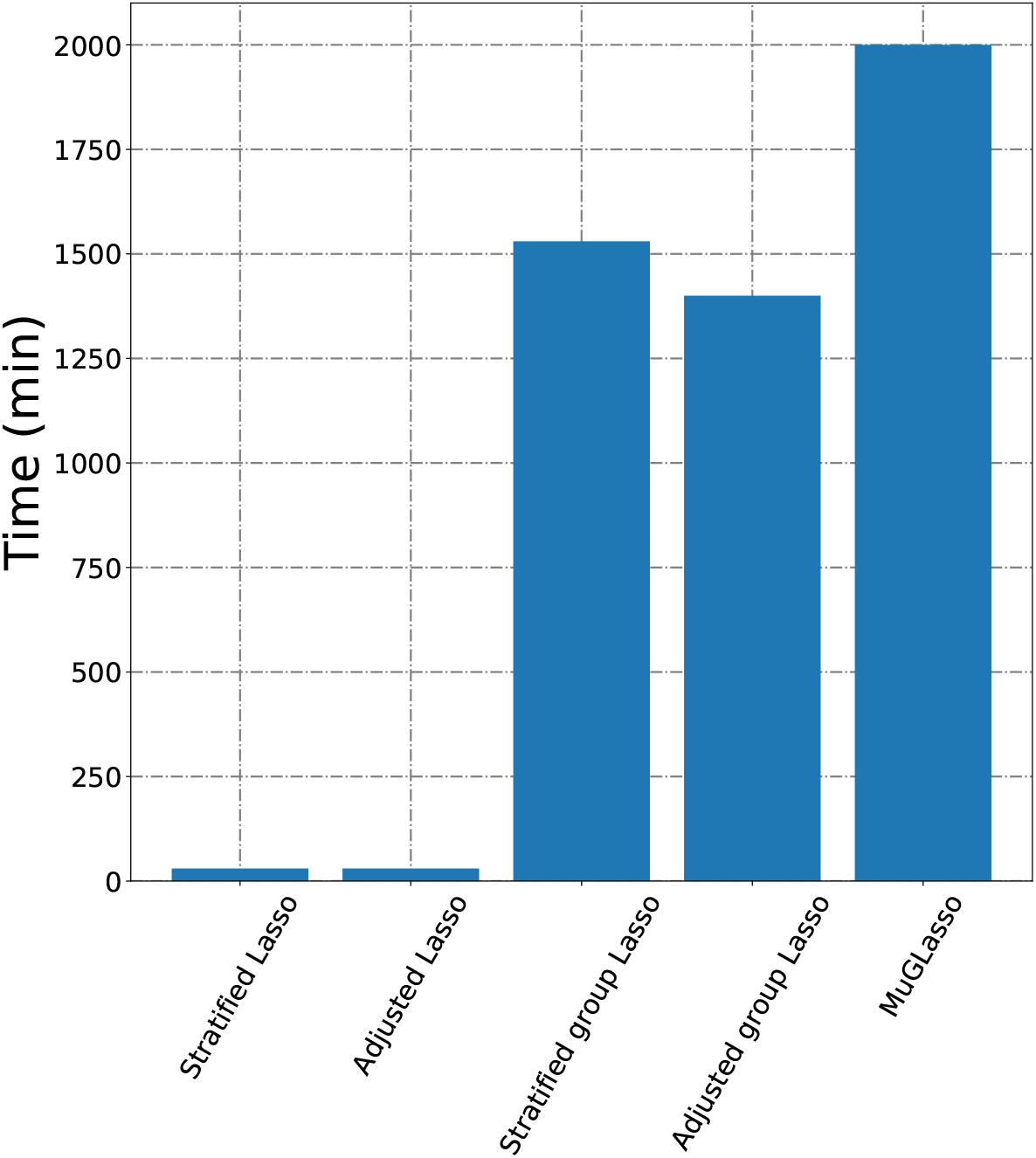
Runtimes of the different Lasso approaches.

#### Appendix C.3. Breast cancer risk loci detected by MuGLasso on DRIVE

On the DRIVE dataset, MuGLasso selected 1 357 SNPs, forming 62 LD groups. Those SNPs include all the 306 SNPs that are significant in the adjusted GWAS approach. We used FUMA [WTVBP17] to analyze the remaining 1051 SNPs, and found that 57% of these SNPs are within 10kb of protein coding genes. Hence MuGLasso identifies a total of 32 genes (listed in in Table C1), in addition to the 9 genes (*ITPR1, MRPS30, MAP3K1, SETD9, MIER3, EBF1, FGFR2, TOX3* and *MKL1*) identified by the adjusted GWAS.

Out of these 32 genes, 17 were previously identified in breast cancer meta-analyses which data include our 28281 samples from the General Research Use dataset of the DRIVE Breast Cancer OncoArray Genotypes (see Table C1). More specifically, these studies respectively used 10707 ER-negative breast cancer cases 76649 controls [GC^+^13] 45290 cases and 41880 controls of European ancestry [M^+^13], 62623 breast cancer cases and 61696 controls [M^+^15], 122 977 cases and 105 974 controls of European ancestry together with 14068 cases and 13104 controls of East Asian ancestry [M^+^17a], and 210 088 controls (9494 of which are BRCA1 mutation carriers) and 30882 cases (21468 ER-negative cases and 9414 BRCA1 mutation carriers), all of European origin [M^+^17].

This suggests that MuGLasso was able to rescue loci that are significant in a better-powered study (that is to say, a study with a larger number of samples).

In addition, we were able to find in the literature prior evidence of relationship with breast cancer risk or tumor growth for 7 additional genes, suggesting biological relevance of the MuGLasso findings.

Further analyses would be required to really get to the biological interpretation of these results. In particular, we restricted ourselves to mapping SNPs to genes based on a 10kb window, where other authors rather use 50kb, and FUMA provides many additional possibilities using known eQTLs and chromatin interactions across all tissues or for relevant tissues. In addition, pathway enrichment analyses could also be very relevant. One could also compare the selected SNPs to those significant in large meta-analyses such as [M^+^17, M^+^17a] in a more systematic manner to investigate how much power is gained by using MuGLasso on a subset of these GWAS data sets. Finally, we have analyzed jointly all selected SNPs and have not distinguished between those that are specific to one of the two populations and those that are common to both.

**Table C1.** The 32 potential breast cancer risk genes within 10kb of loci identified by MuGLasso and not the adjusted GWAS, together with information as to their biological relevance.

| Genes found in meta-GWAS including the samples used in this work |  |
| --- | --- |
| Gene symbols | Evidence |
| *ASTN2* | M<sup>+</sup>17a |
| *CCDC170* | GC<sup>+</sup>13,M<sup>+</sup>13,M<sup>+</sup>15,M<sup>+</sup>17a,M<sup>+</sup>17 |
| *CDYL2* | M<sup>+</sup>13,M<sup>+</sup>15,M<sup>+</sup>17a |
| *DIRC3* | M<sup>+</sup>13,M<sup>+</sup>15,M<sup>+</sup>17a,M<sup>+</sup>17 |
| *ELL* | M<sup>+</sup>13,M<sup>+</sup>15,M<sup>+</sup>17a,M<sup>+</sup>17 |
| *ESR1* | GC<sup>+</sup>13,M<sup>+</sup>15,M<sup>+</sup>17a,M<sup>+</sup>17 |
| *FTO* | GC<sup>+</sup>13,M<sup>+</sup>13,M<sup>+</sup>15,M<sup>+</sup>17a,M<sup>+</sup>17 |
| *GRHL1* | M<sup>+</sup>17a |
| *KCNU1* | M<sup>+</sup>15,M<sup>+</sup>17a |
| *NEK10* | M<sup>+</sup>13,M<sup>+</sup>15,M<sup>+</sup>17a,M<sup>+</sup>17 |
| *PAX9* | M<sup>+</sup>13,M<sup>+</sup>15,M<sup>+</sup>17a |
| *PTHLH* | GC<sup>+</sup>13,M<sup>+</sup>13,M<sup>+</sup>15,M<sup>+</sup>17a,M<sup>+</sup>17 |
| *SSBP4* | M<sup>+</sup>17a |
| *TGFBR2* | M<sup>+</sup>13,M<sup>+</sup>15,M<sup>+</sup>17a |
| *TNRC6B* | M<sup>+</sup>17a |
| *ZMIZ1* | M<sup>+</sup>13,M<sup>+</sup>15,M<sup>+</sup>17a |
| *ZNF365* | M<sup>+</sup>17a,M<sup>+</sup>17 |
| Genes found to be associated with breast cancer risk or tumor growth in the literature |  |
| Gene symbols | Evidence |
| *ADSL* | oncogenic driver in triple negative breast cancer [Z<sup>+</sup>19] |
| *CACNA1I* | underexpressed in breast cancer [P<sup>+</sup>17] |
| *CCDC91* | likely target gene of breast cancer risk variants [F<sup>+</sup>19] |
| *NUP205* | forms a complex with NUP93 which regulates breast tumor growth [B<sup>+</sup>20] |
| *POP1* | expression correlates with prognosis in breast cancer [L<sup>+</sup>21] |
| *PPFIBP1* | promotes cell motility and migration in breast cancer [C<sup>+</sup>16] |
| *SGSM3* | associated with breast cancer in a Chinese population [TZS17] |
| Other genes |  |
| *C7orf73*, *CCSER1*, *CD2AP*, *HK1*, *HRSP12*, *LUC7L3*, *MED21*, *REP15* |  |

## Footnotes

a https://github.com/EugeneNdiaye/Gap_Safe_Rules

